# Lighter: fast and memory-efficient error correction without counting

**DOI:** 10.1101/005579

**Authors:** Li Song, Liliana Florea, Ben Langmead

## Abstract

Lighter is a fast, memory-efficient tool for correcting sequencing errors. Lighter avoids counting *k*-mers. Instead, it uses a pair of Bloom filters, one holding a sample of the input *k*-mers and the other holding *k*-mers likely to be correct. As long as the sampling fraction is adjusted in inverse proportion to the depth of sequencing, Bloom filter size can be held constant while maintaining near-constant accuracy. Lighter is parallelized, uses no secondary storage, and is both faster and more memory-efficient than competing approaches while achieving comparable accuracy.

## Introduction

The cost and throughput of DNA sequencing have improved rapidly in the past several years [1], with recent advances reducing the cost of sequencing a single human genome at 30-fold coverage to around $1,000 [2]. With these advances has come an explosion of new software for analyzing large sequencing datasets. Sequencing error correction is a basic need for many of these tools. Removing errors at the outset of an analysis can improve accuracy of downstream tools such as variant callers [3]. Removing errors can also improve the speed and memory-efficiency of downstream tools, particularly for de novo assemblers based on De Bruijn graphs [4, 5].

To be useful in practice, error correction software must make economical use of time and memory even when input datasets are large (many billions of reads) and when the genome under study is also large (billions of nucleotides). Several methods have been proposed, covering a wide tradeoff space between accuracy, speed and memory- and storage-efficiency. SHREC [6] and HiTEC [7] build a suffix index of the input reads and locate errors by finding instances where a substring is followed by a character less often than expected. Coral [8] and ECHO [9] find overlaps among reads and use the resulting multiple alignments to detect and correct errors. Reptile [10] and Hammer [11] detect and correct errors by examining each *k*-mer’s neighborhood in the dataset’s *k*-mer Hamming graph.

The most practical and widely used error correction methods descend from the spectral alignment approach introduced in the earliest De Bruijn graph based assemblers [4, 5]. These methods count the number of times each *k*-mer occurs (its *multiplicity)* in the input reads, then apply a threshold such that reads with multiplicity exceeding the threshold are considered *solid*. These *k*-mers are unlikely to have been altered by sequencing errors. *k*-mers with low multiplicity (*weak k*-mers) are systematically edited into high-multiplicity *k*-mers using a dynamic-programming solution to the spectral alignment problem [4, 5] or, more often, a fast heuristic approximation. Quake [3], the most widely used error correction tool, uses a hash-based *k*-mer counter called Jellyfish [12] to determine which *k*-mers are correct. CUDA-EC [13] was the first to use a Bloom filter as a space-efficient alternative to hash tables for counting *k*-mers and for representing the set of solid *k*-mers. More recent tools such as Musket [14] and BLESS [15] use a combination of Bloom filters and hash tables to count *k*-mers or to represent the set of solid *k*-mers.

*Lighter* (LIGHTweight ERror corrector) is also in the family of spectral alignment methods, but differs from previous approaches in that it avoids counting *k*-mers. Rather than count *k*-mers, Lighter samples *k*-mers randomly, storing the sample in a Bloom filter. Lighter then uses a simple test applied to each position of each read to compile a set of solid *k*-mers, stored in a second Bloom filter. These two Bloom filters are the only sizable data structures used by Lighter.

A crucial advantage is that Lighter’s parameters can be set such that memory footprint and accuracy are near-constant with respect to depth of sequencing. That is, no matter how deep the coverage, Lighter can allocate the same sized Bloom filters and achieve nearly the same (a) Bloom filter occupancy, (b) Bloom filter false positive rate, and (c) error correction accuracy. Lighter does this without using any disk space or other secondary memory. This is in contrast to BLESS and Quake/Jellyfish, which use secondary memory to store some or all of the *k*-mer counts.

Lighter’s accuracy is comparable to competing tools. We show this both in simulation experiments where false positives and false negatives can be measured, and in real-world experiments where read alignment scores and assembly statistics can be measured. Lighter is also very simple and fast, faster than all other tools tried in our experiments. These advantages make Lighter quite practical compared to previous counting-based approaches, all of which require an amount of memory or secondary storage that increases with depth of coverage. Lighter is free open source software available from https://github.com/mourisl/Lighter/.

## Method

Lighter’s workflow is illustrated in Figure 1. Lighter makes three passes over the input reads. The first pass obtains a sample of the *k*-mers present in the input reads, storing the sample in Bloom filter A. The second pass uses Bloom filter A to identify solid *k*-mers, which it stores in Bloom filter B. The third pass uses Bloom filter B and a greedy procedure to correct errors in the input reads.

**Figure 1.**
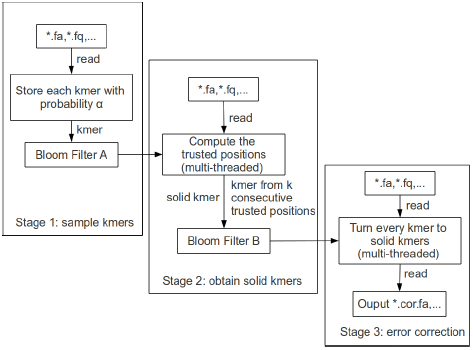
The framework of Lighter.

### Bloom filter

A Bloom filter [16] is a compact probabilistic data structure representing a set. It consists of an array of *m* bits, each initialized to 0. To add an item *o, h* independent hash functions *H*_0_(*o*)*, H*_1_(*o*)*, …, H*_h-1_(*o*) are calculated. Each maps *o* to an integer in [0*, m*) and the corresponding *h* array bits are set to 1. To test if item *q* is a member, the same hash functions are applied to *q*. *q* is a member if all corresponding bits are set to 1. A false positive occurs when the corresponding bits are set to 1 “by coincidence,” that is, because of items besides *q* that were added previously. Assuming the hash functions map items to bit array elements with equal probability, the Bloom filter’s false positive rate is approximately 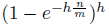, where *n* is the number of distinct items added, which we call the *cardinality*. Given *n*, which is usually determined by the dataset, *m* and *h* can be adjusted to achieve a desired false positive rate. Lower false positive rates can come at a cost, since greater values of *m* require more memory and greater values of *h* require more hash function calculations. Many variations on Bloom filters have been proposed that additionally permit compression of the filter, storage of count data, representation of maps in addition to sets, etc [17]. Bloom filters and variants thereon have been applied in various bioinformatics settings, including assembly [18], compression [19], *k*-mer counting [20], and error correction [13].

By way of contrast, another way to represent a set is with a hash table. Hash tables do not yield false positives, but Bloom filters are far smaller. Whereas a Bloom filter is an array of bits, a hash table is an array of buckets, each large enough to store a pointer, key, or both. If chaining is used, lists associated with buckets incur additional overhead. While the Bloom filter’s small size comes at the expense of false positives, these can be tolerated in many settings including in error correction.

Lighter’s efficiency depends on the efficiency of the Bloom filter implementation. Specifically Lighter uses a *pattern-blocked* Bloom filter to decrease overall number of cache misses and improve efficiency. This comes at the expense of needing a slightly larger filter to achieve a comparable false positive rate to a standard filter, as discussed in Supplementary Note 1.

In our method, the items to be stored in the Bloom filters are *k*-mers. Because we would like to treat genome strands equivalently for counting purposes, we will always *canonicalize* a *k*-mer before adding it to, or using it to query a Bloom filter. A canonicalized *k*-mer is either the *k*-mer itself or its reverse complement, whichever is lexicographically prior.

### Sequencing model

We use a simple model to describe the sequencing process and Lighter’s subsampling. The model resembles one suggested previously [21]. Let *K* be the total number of *k*-mers obtained by the sequencer. We say a *k*-mer is *incorrect* if its sequence has been altered by one or more sequencing errors. Otherwise it is *correct*. Let *ϵ* be the fraction of *k*-mers that are incorrect. We assume *ϵ* does not vary with the depth of sequencing. The sequencer obtains correct *k*-mers by sampling independently and uniformly from *k*-mers in the genome. Let the number of *k*-mers in the genome be *G*, and assume all are distinct. If *κ*_*c*_ is a random variable for the multiplicity of a correct *k*-mer in the input, *κ*_*c*_ is binomial with success probability 1*/G* and number of trials (1 − *ϵ*)*K*: *κ*_*c*_ ∼ *Binom*((1 - *ϵ*)*K,* 1*/G*). Since the number of trials is large and the success probability is small, the binomial is well approximated by a Poisson: *κ*_*c*_ ∼ *Pois*(*K*(1 - *ϵ*)*/G*)

A sequenced *k*-mer survives subsampling with probability *α*. If 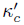 is a random variable for the number of times a correct *k*-mer appears in the subsample, 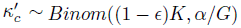, which is approximately *Pois*(*αK*(1 - *ϵ*)*/G*).

We model incorrect *k*-mers similarly. The sequencer obtains incorrect *k*-mers by sampling independently and uniformly from *k*-mers “close to” a *k*-mer in the genome. We might define these as the set of all *k*-mers with low but non-zero Hamming distance from some genomic *k*-mer. If *κ*_*e*_ is a random variable for the multiplicity of an incorrect *k*-mer, *κ*_*e*_ is binomial with success probability 1*/H* and number of trials *ϵK*: *κ*_*e*_ ∼ *Binom*(*ϵK,* 1*/H*), which is approximately *Pois*(*Kϵ/H*). It is safe to assume 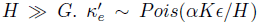 is a random variable for the number of times an incorrect *k*-mer appears in the subsample.

Others have noted that, given a dataset with deep and uniform coverage, incorrect *k*-mers occur rarely while correct *k*-mers occur many times, proportionally to coverage [4, 5].

### Stages of the method

#### First pass

In the first pass, Lighter examines each *k*-mer of each read. With probability 1 - *α*, the *k*-mer is ignored. *k*-mers containing ambiguous nucleotides (e.g. “N”) are also ignored. Otherwise, the *k*-mer is canonicalized and added to Bloom filter *A*.

Say a distinct *k*-mer *a* occurs a total of *N*_*a*_ times in the dataset. If none of the *N*_*a*_occurrences survive subsampling, the *k*-mer is never added to *A* and *A*’s cardinality is reduced by one. Thus, reducing *α* can in turn reduce *A*’s cardinality. Because correct *k*-mers are more numerous, incorrect *k*-mers tend to be discarded from *A* before correct *k*-mers as *α* decreases.

The subsampling fraction *α* is set by the user. We suggest adjusting *α* in inverse proportion to depth of sequencing, for reasons discussed below. For experiments described here, we set *α* = 0.1 when the average coverage is 70-fold. That is, we set *α* to 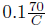 where *C* is average coverage.

#### Second pass

A read position is overlapped by up to *x k*-mers, 1 ≤ *x* ≤ *k*, where *x* depends on how close the position is to either end of the read. For a position altered by sequencing error, the overlapping *k*-mers are all incorrect and are unlikely to appear in *A*. We apply a threshold such that if the number of *k*-mers overlapping the position and appearing in Bloom filter *A* is less than the threshold, we say the position is *untrusted*. Otherwise we say it is *trusted*. Each instance where the threshold is applied is called a *test case*. When one or more of the *x k*-mers involved in two test cases differ, we say the test cases are distinct.

Let *P* ^∗^(*α*) be the probability an incorrect *k*-mer appears in *A*, taking the Bloom filter’s false positive rate into account. If random variable *B*_*e,x*_ represents the number of *k*-mers appearing in *A* for an untrusted position overlapped by *x k*-mers, *B*_*e,x*_ ∼ *Binom*(*x, P* ^∗^(*α*)). We define thresholds *y*_*x*_, for each *x* in [1*, k*]. *y*_*x*_ is the minimum integer such that *p*(*B*_*e,x*_ ≤ *y*_*x*_ − 1) ≥ 0.995.

Ignoring false positives for now, we model the probability of a sequenced a *k*-mer having been added to *A* as *P* (*α*) = 1−(1−*α*)^*f*(*α*)^. We define *f*(*α*) = *max*{2, 0.2*/α*}. That is, we assume the multiplicity of a weak *k*-mer is at most *f*(*α*), which will often be a conservative assumption, especially for small *α*. It is also possible to define *P*(*α*) in terms of random variables *κ*_*e*_ and 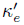, but we avoid this here for simplicity.

A property of this threshold is that when *α* is small, *P* (*α/z*) = 1−(1−*α/z*)^0.2z/α^ ≈ 1 − (1 − *α*)^0.2/α^ = *P* (*α*), where *z* is a constant greater than 1 and we use the fact that (1 − *α/z*)^z^ ≈ 1 − *α*.

For *P* ^∗^(*α*), we additionally take *A*’s false positive rate into account. If the false positive rate is *β*, then *P* ^∗^(*α*) = *P* (*α*) + *β* − *βP* (*α*).

Once all positions in a read have been marked *trusted* or *untrusted* using the threshold, we find all instances where *k* trusted positions appear consecutively. The *k*-mer made up by those positions is added to Bloom filter *B*.

#### Third pass

In the third pass, Lighter applies a simple, greedy error correction procedure similar to that used in BLESS [15]. A read *r* of length |*r*|, contains |*r*| − *k* + 1 *k*-mers. *k*_*i*_ denotes the *k*-mer starting at read position *i*, 1 ≤ *i* ≤ |*r*| − *k* + 1. We first identify the longest stretch of consecutive *k*-mers in the read that appear in Bloom filter *B*. Let *k*_*b*_ and *k*_*e*_ be the *k*-mers at the left and right extremes of the stretch. If *e <* |*r*| − *k* + 1, we examine successive *k*-mers to the right starting at *k*_*e*_ + 1. For a *k*-mer *k*_*i*_ that does not appear in *B*, we assume the nucleotide at offset *i* + *k* − 1 is incorrect. We consider all possible ways of substituting for the incorrect nucleotide. For each substitution, we count how many consecutive *k*-mers starting with *k*_*i*_ appear in Bloom filter *B* after making the substitution. We pick the substitution that creates the longest stretch of consecutive *k*-mers in *B*. The procedure is illustrated in Figure 2.

**Figure 2.**
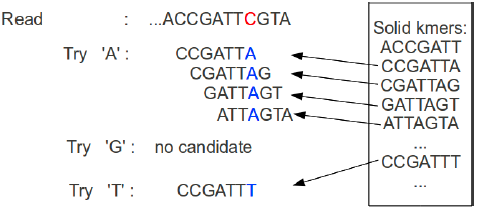
An example of the greedy error correction procedure. *k*-mer CCGATTC does not appear in Bloom filter *B*, so we attempt to substitute a different nucleotide for the C shown in red. We select A since it yields the longest stretch of consecutive *k*-mers that appear in Bloom filter *B*.

If more than one candidate substitution is equally good (i.e. results in the same number of consecutive *k*-mers from *B*), we call position *i* + *k* − 1 ambiguous and make no attempt to correct it. The procedure then resumes starting at *k*_i+k_, or the procedure ends if the read is too short to contain *k*-mer *k*_*i+k*_.

When errors are located near to end of a read, the stretches of consecutive *k*-mers used to prioritize substitutions are short. E.g. if the error is at the very last position of the read, we must choose a substation on the basis of just one *k*-mer: the rightmost *k*-mer. This very often results in a tie, and no correction. Lighter avoid many of these ties by considering *k*-mers that extend beyond the end of the read, as discussed in Supplementary Note 2.

For better precision, Lighter also limits the corrections that can be made in any window of size *k* in a read. The default limit is 4, and it is configurable. Corrections at positions with an “N” contribute 0, and corrections at low-quality bases (defined in the Quality score section below) contribute 0.5 toward this limit. All other positions contribute 1.

### Scaling with depth of sequencing

Lighter’s accuracy can be made near-constant as the depth of sequencing *K* increases and its memory footprint is held constant. This is accomplished by holding *αK* constant, i.e., by adjusting *α* in inverse proportion to *K*. This is illustrated in Tables 1 and 2. We also argue this more formally in Supplementary Note 3.

**Table 1.** Accuracy measures for datasets simulated with Mason with various sequencing depths and error rates. Rows labeled *k* show the *k*-mer sizes selected for each tool and dataset.

| Coverage |  | 35× |  | 70× |  | 140× |  |
| --- | --- | --- | --- | --- | --- | --- | --- |
| Error rate |  | 1% | 3% | 1% | 3% | 1% | 3% |
| $\alpha$ for Lighter | | 0.2 | 0.2 | 0.1 | 0.1 | 0.05 | 0.05 |
| Recall | Quake | 89.68 | 48.77 | 89.64 | 48.82 | 89.59 | 48.78 |
|  | SOAPec | 57.71 | 38.00 | 57.57 | 37.71 | 57.09 | 36.76 |
|  | Musket | 93.75 | 92.62 | 93.73 | 92.64 | 93.73 | 92.63 |
|  | Bless | 99.81 | **99.33** | 99.82 | **99.58** | 99.82 | **99.58** |
|  | Lighter | **99.87** | 98.53 | **99.84** | 98.72 | **99.86** | 98.78 |
| Prec | Quake | **99.99** | 99.99 | **99.99** | **99.99** | **99.99** | **99.99** |
|  | SOAPec | **99.99** | **100.00** | **99.99** | **99.99** | **99.99** | **99.99** |
|  | Musket | **99.99** | 99.93 | **99.99** | 99.93 | **99.99** | 99.93 |
|  | Bless | 99.73 | 98.86 | 99.73 | 99.35 | 99.72 | 99.36 |
|  | lighter | 99.98 | 99.96 | 99.98 | 99.96 | 99.98 | 99.96 |
| F-score | Quake | 94.55 | 65.56 | 94.54 | 65.61 | 94.51 | 65.57 |
|  | SOAPec | 73.18 | 55.07 | 73.07 | 54.77 | 72.68 | 53.75 |
|  | Musket | 96.77 | 96.14 | 96.76 | 96.15 | 96.76 | 96.15 |
|  | Bless | 99.77 | 99.09 | 99.77 | **99.47** | 99.77 | **99.47** |
|  | Lighter | **99.93** | **99.24** | **99.91** | 99.33 | **99.92** | 99.36 |
| Gain | Quake | 89.67 | 48.76 | 89.64 | 48.82 | 89.59 | 48.78 |
|  | SOAPec | 57.70 | 38.00 | 57.57 | 37.71 | 57.09 | 36.75 |
|  | Musket | 93.74 | 92.56 | 93.72 | 92.58 | 93.72 | 92.57 |
|  | Bless | 99.54 | 98.19 | 99.54 | **98.93** | 99.54 | **98.94** |
|  | Lighter | **99.85** | **98.49** | **99.81** | 98.68 | **99.84** | 98.73 |
| $k$ | Quake | 17 | 17 | 17 | 17 | 17 | 17 |
|  | SOAPec | 17 | 17 | 17 | 17 | 17 | 17 |
|  | Musket | 23 | 19 | 23 | 19 | 23 | 19 |
|  | Bless | 31 | 23 | 31 | 23 | 31 | 23 |
|  | Lighter | 23 | 19 | 23 | 19 | 23 | 19 |

**Table 2.**
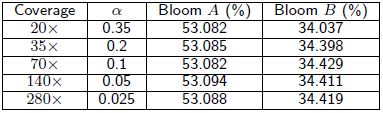
Occupancy (fraction of bits set) for Bloom filters *A* and *B* for various coverages

| Coverage | $\alpha$ | Bloom $A$ (%) | Bloom $B$ (%) |
| --- | --- | --- | --- |
| 20× | 0.35 | 53.082 | 34.037 |
| 35× | 0.2 | 53.085 | 34.398 |
| 70× | 0.1 | 53.082 | 34.429 |
| 140× | 0.05 | 53.094 | 34.411 |
| 280× | 0.025 | 53.088 | 34.419 |

### Quality score

A low base quality value at a certain position can force Lighter to treat that position as untrusted even if the overlapping *k*-mers indicate it is trusted. First, Lighter scans the first 1 million reads in the input, recording the quality value at the last position in each read. Lighter then chooses the 5th-percentile quality value; that is, the value such that 5% of the values are less than or equal to it say *t*_1_. Use the same idea, we get another 5th-percentile quality, say *t*_2_ value for the first 1 million reads’ first base. When Lighter decides whether a position is trusted or not, if its quality score is less or equal to *min*{*t*_1_*, t*_2_ - 1}, then call it untrusted regardless of how many of the overlapping *k*-mers appear in Bloom filter *A*.

### Parallelization

As shown in Figure 1, Lighter works in three passes: (1) populating Bloom filter *A* with a *k*-mer subsample, (2) applying the per-position test and populating Bloom filter *B* with likely-correct *k*-mers, and (3) error correction. For pass 1, because *α* is usually small, most time is spent scanning the input reads. Consequently, we found little benefit to parallelizing pass 1. Pass 2 is parallelized by using concurrent threads handle subsets of input reads. Because Bloom filter *A* is only being queried (not added to), we need not synchronize accesses to *A*. Accesses to *B* are synchronized so that additions of *k*-mers to *B* by different threads do not interfere. Since it is typical for the same correct *k*-mer to be added repeatedly to *B*, we can save synchronization effort by first checking whether the *k*-mer is already present and adding it (synchronously) only if necessary. Pass 3 is parallelized by using concurrent threads to handle subsets of the reads; since Bloom filter *B* is only being queried, we need not synchronize accesses.

## Evaluation

Supplementary Note 4 describes the computer all experiments were conducted on. Supplementary Note 5 describes the exact command lines used.

### Simulated dataset

#### Accuracy on simulated data

We compared Lighter v1.0.2’s performance with Quake v0.3 [3], Musket v1.1 [14], BLESS v0p17 [15], and SOAPec v2.0.1 [22]. We simulated a collection of reads from the reference genome for the K12 strain of *E. coli* (NC 000913.2) using Mason v0.1.2 [23].

We simulated six distinct datasets with 101 bp single-end reads, varying average coverage (35x, 75x 140x) and average error rate (1% and 3%). For a given error rate *e* we specify Mason parameters -qmb *e/*2 -qme 3*e*, so that the average error rate is *e* but errors are more common toward the 3’ end, as in real datasets.

We then ran all four tools on all six datasets, with results presented in Table 1. BLESS was run with the `-notrim` option to make the results more comparable. In these comparisons, a true positive (TP) is an instance where an error is successfully corrected, i.e. with the correct base substituted. A false positive (FP) is an instance where a spurious substitution is made at an error-free position. A false negative (FN) is an instance where we either fail to detect an error or an incorrect base is substituted. As done in previous studies [14], we report the following summaries: recall = TP/(TP + NP), precision = TP/(TP + FP), F-score = 2×recall×precision/(recall + precision) and gain = (TP-FP)/(TP + FN).

Since these tools are sensitive to the choice of *k*-mer size, we tried several values for this parameter (17, 19, 23, 27, 31) and picked the value yielding the greatest gain in the accuracy evaluation. The *k*-mer sizes chosen are shown in the bottom rows of Table 1. Note that SOAPec’s maximum *k*-mer size is 27. We found that Quake crashed for *k*-mer sizes 23 and up.

Unlike the other tools, Quake both trims the untrusted tails of the reads and discards reads it cannot correct. BLESS also trims some reads (even in `-notrim` mode), but only a small fraction (*<*0.1%) of them, which has only a slight effect on results. For these simulation experiments, we measure precision and recall with respect to all the nucleotides (even the trimmed ones) in all the reads (even those discarded). This tends to lead to higher precision but lower recall for Quake relative to the other tools.

Apart from Quake, SOAPec, Musket and Lighter achieve the highest precision. Lighter achieves the highest recall, F-score and gain in the experiments with 1% error, and is comparable to BLESS when the error rate is 3%.

To see how quality value information affects performance, we repeated these experiments with quality values omitted (Supplementary Table 1). Quake and BLESS accept only FASTQ input files (which include quality values), and so were not included in the experiment. Lighter achieves superior recall, gain and F-score.

To see how the choice of read simulator affects performance, we repeated these experiments using the Art [24] simulator to generate the reads instead of Mason (Supplementary Table 2). All tools perform quite similarly in this experiment, except SOAPec, which has poor recall compared to the others. There is less difference between tools than in the Mason experiment, likely because Art simulates a relatively low (∼0.25%) error rate. Lighter and Musket perform best overall.

For the Mason-simulated 1% error dataset, we found that Lighter’s gain was maximized by setting the *k*-mer size to 23. We therefore fix the *k*-mer size to 23 for subsequent experiments, except where otherwise noted.

#### C. elegans simulation

We performed a similar accuracy test as in the previous section, but using data simulated from the larger *C. elegans* genome, WBcel235 (Supplementary Table 3). We used Mason to simulate a dataset of 101 bp single-end reads with a 1% error rate totaling 35x coverage. We again tried several values for the *k*-mer size parameter (19, 23, 27, 31) and picked the value yielding the greatest gain in the accuracy evaluation. As for the *E. coli* experiment, Lighter had the greatest recall, F-score and gain.

#### Scaling with depth of simulated sequencing

We also used Mason to generate a series of datasets with 1% error, similar to those used in Table 1, but for 20×, 35×, 70×, 140× and 280× average coverage. We ran Lighter on each and measured final occupancies (fraction of bits set) for Bloom filters *A* and *B*. If our assumptions and scaling arguments are accurate, we expect the final occupancies of the Bloom filters to remain approximately constant for relatively high levels of coverage. As seen in Table 2, this is indeed the case.

#### Cardinality of Bloom filter B

We also measured the number of correct *k*-mers added to table *B*. We used the Mason dataset with 70x coverage and 1% error rate. The *E. coli* genome has 4,564,614 distinct *k*-mers, and 4,564,569 (99.999%) of them are in table *B*.

#### Effect of ploidy on Bloom filter B

We conducted a experiment similar to that in the previous section but with Mason configured to simulate reads from a diploid version of the *E. coli* genome. Specifically, we introduced heterozygous SNPs at 0.1% of the positions in the reference genome. Mason then sampled equal numbers of reads from both genomes, making a dataset with 70x average coverage in total. Of the 214,567 simulated *k*-mers that overlapped a position with a heterozygous SNP, table *B* held 214,545 (99.990%) of them at the end of the run. Thus, Lighter retained in table B almost the same fraction of the *k*-mers overlapping heterozygous positions (99.990%) as of the *k*-mers overall (99.999%).

Musket and BLESS both infer a threshold for the multiplicity of solid *k*-mers. In this experiment, Musket inferred a threshold of 10 and BLESS inferred a threshold of 9. All three tools are using a *k*-mer size of 23. By counting the multiplicity of the *k*-mers overlapping heterozygous positions, we conclude that Musket would classify 214,458 (99.949%) as solid and BLESS would classify 214,557 (99.995%) as solid. So in the diploid case, it seems Lighter’s ability to identify correct *k*-mers overlapping heterozygous SNPs is comparable to that of error correctors that are based on counting.

Diploidy is one example of a phenomenon that tends to drive the count distribution for some correct *k*-mers (those overlapping heterozygous variants) closer to the count distribution for incorrect *k*-mers. In the Discussion section we elaborate on other such phenomena, such as copy number, sequencing bias, and non-uniform coverage.

#### Effect of varying α

In a series of experiments, we measured how different settings for the subsampling fraction *α* affected Lighter’s accuracy as well as the occupancies of Bloom filters *A* and *B*. We still use the datasets simulated by Mason with 35×, 70× and 140× coverage.

As shown in Figures 3 and 4, only a fraction of the correct *k*-mers are added to *A* when *α* is very small, causing many correct read positions to fail the threshold test. Lighter attempts to “correct” these error-free positions, decreasing accuracy. This also has the effect of reducing the number of consecutive stretches of *k* trusted positions in the reads, leading to a smaller fraction of correct *k*-mers added to *B*, and ultimately to lower accuracy. When *α* grows too large, the *y*_*x*_ thresholds grow to be greater than *k*, causing all positions to fail the threshold test, as seen in Figure 4’s right-hand side. This also leads to a dramatic drop in accuracy as seen in Figure 3. Between the two extremes, we find a fairly broad range of values for *α* (from about 0.15 to 0.3) that yield high accuracy when the error rate is 1% or 3%. The range is wider when the error rate is lower.

**Figure 3.**
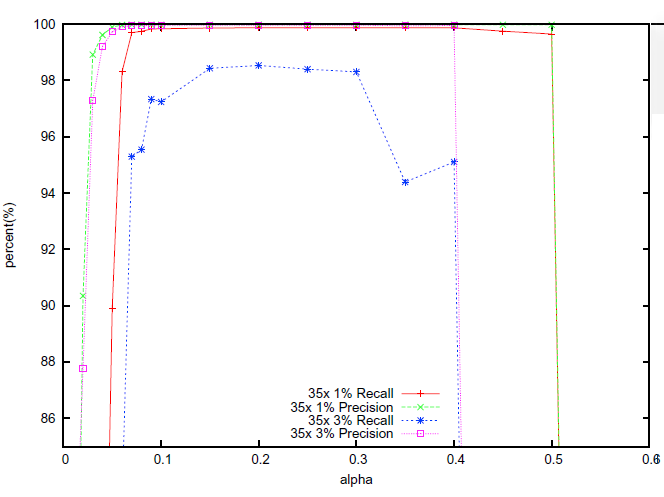
The effect of *α* on the accuracy using the simulated 35× dataset.

**Figure 4.**
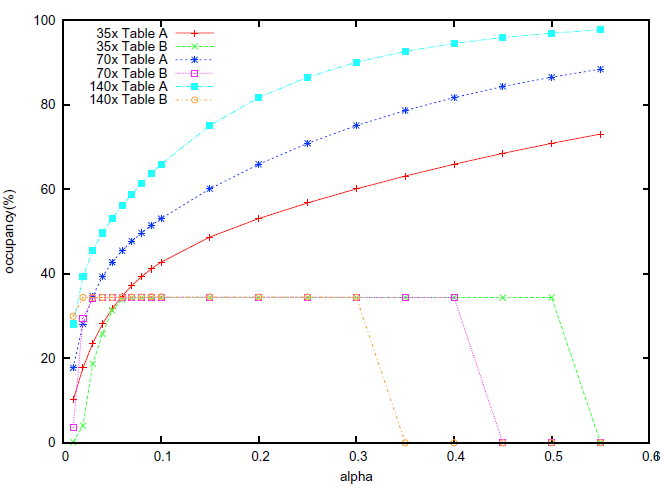
The effect of *α* on occupancy of Bloom filters *A* and *B*. The effect of α on occupancy of Bloom filters A and B using simulated 35×, 70× and 140× datasets. The error rate is 1%

#### Effect of varying k

A key parameter of Lighter is the *k*-mer length *k*. Smaller *k* yields higher probability that a *k*-mer affected by a sequencing error also appears elsewhere in the genome. For larger *k*, the fraction of *k*-mers that are correct decreases, which could lead to fewer correct *k*-mer in Bloom filter *A*. We measured how different settings for *k* affect accuracy using the simulated data with 35× coverage and both 1%, 3% error rate. Results are shown in Figure 5. Accuracy is high for *k*-mer lengths ranging from about 18 to 30 when the error rate is 1%. But the recall drops gradually when when the rror rate is 3%.

**Figure 5.**
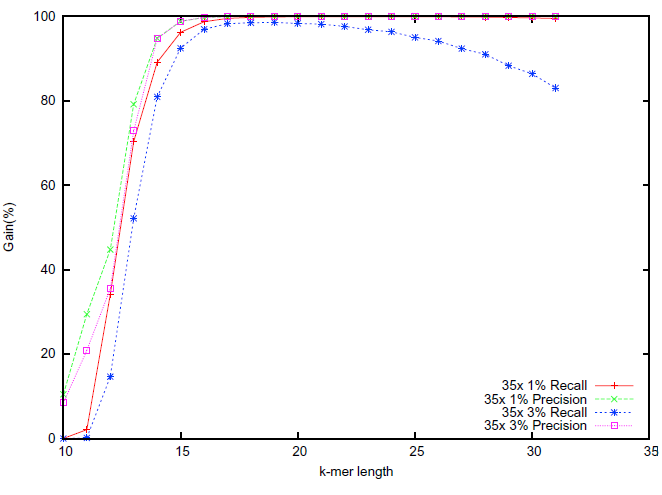
The effect of *k*-mer length *k* on accuracy.

### Real datasets

*E. coli.* Next we benchmarked the same error correction tools using a real sequencing dataset, ERR022075. This is a deep DNA sequencing dataset of the the K-12 strain of the *E. coli* genome. To obtain a level of coverage more reflective of other projects, we randomly subsampled the reads in the dataset to obtain roughly 75x coverage (∼3.5M reads) of the *E. coli* K-12 reference genome. The reads are 100 × 102 bp paired-end reads. Because BLESS cannot handle paired-end reads where the ends have different lengths, we truncated the last 2 bases from the 102 bp end before running our experiments. We again ran BLESS with the `-notrim` option.

These data are not simulated, so we cannot measure accuracy directly. But we can measure it indirectly, as other studies have [15], by measuring read alignment statistics before and after error correction. We use Bowtie2 [25] v2.2.2 with default parameters to align the original reads and the corrected reads to the *E. coli* K-12 reference genome. For each error corrector, we tested different *k*-mer sizes (17, 19, 23, 27, 31) and chose the size that yielded the greatest total number of matching aligned nucleotides. For Quake and BLESS, we use only the reads (and partial reads) that remained after trimming and discarding for this evaluation. Results are shown in Table 3. Lighter yields the greatest improvement in fraction of reads aligned, whereas Quake and BLESS yield the greatest improvement in fraction of aligned bases that match the reference, with Lighter very close behind. As before, Quake is hard to compare to the other tools because it trims and discards many reads.

**Table 3.** Alignment statistics for the 75× *E. coli* dataset, before error correction (Original) and after error correction (other rows). k column shows k-mer size selected for each tool. First “Increase” column shows percent increase in reads aligned. Second “Increase” column shows percent increase in the fraction of aligned bases that match the reference genome.

| | $k$ | Read Level | | Base Level | |
| --- | --- | --- | --- | --- | --- |
|  |  | Mapped Reads | Increase (%) | Matches/aligned base (%) | Increase (%) |
| Original | - | 3,464,137 | - | 99.038 | - |
| Quake | 19 | 3,373,498 | -2.62 | 99.659 | 0.63 |
| SOAPec | 17 | 3,465,819 | 0.05 | 99.130 | 0.09 |
| Musket | 17 | 3,467,875 | 0.11 | 99.601 | 0.57 |
| BLESS | 19 | 3,468,677 | 0.13 | 99.666 | 0.63 |
| Lighter | 19 | 3,478,658 | 0.42 | 99.639 | 0.61 |

We repeated this experiment using a less sensitive setting for Bowtie 2 (Supplementary Table 4) and using BWA-MEM [26] v0.7.9a-r786 to align the reads instead of Bowtie 2 (Supplementary Table 5) and found that the error correction tools performed similarly relative to each other.

**Table 4.** De novo assembly statistics for the *E. coli* dataset

|  | N50 | NG50 | Edits / 100kbp | Misassemblies | Coverage (%) |
| --- | --- | --- | --- | --- | --- |
| Original | 94,879 | 94,879 | 3.41 | 0 | 97.496 |
| Quake | 89,470 | 88,209 | 11.62 | 4 | 97.515 |
| SOAPec | 98,111 | 94,879 | 3.49 | 1 | 97.473 |
| Musket | 86,421 | 86,421 | 6.45 | 0 | 97.53 |
| BLESS | 85,486 | 85,486 | 3.58 | 1 | 97.302 |
| Lighter | 105,460 | 105,460 | 3.71 | 1 | 97.477 |

To further assess accuracy, we assembled the reads before and after error correction and measured relevant assembly statistics using Quast [27]. The corrected reads are those reported in Table 3. We used Velvet 1.2.10 [28] to assemble. Velvet is a De Bruijn graph-based assembler designed for second-generation sequencing reads. A key parameter of Velvet is the De Bruijn graph’s *k*-mer length. For each tool we tested different *k*-mer sizes for Velvet (43, 47, 49, 51, 53, 55, 57, 63, 67) and chose the one that yielded the greatest NG50. We set the error correctors’ *k*-mer sizes to match those selected in the alignment experiment of Table 3. As before, we used only the reads (and partial reads) that remained after trimming and discarding for Quake and BLESS. For each assembly, we then evaluated the assembly’s quality using Quast, which was configured to discard contigs shorter than 100 bp before calculating statistics. Results are shown in Table 4.

N50 is the length such that the total length of the contigs no shorter than the N50 cover at least half the assembled genome. NG50 is similar, but with the requirement that contigs cover half the reference genome rather than half the assembled genome. Edits per 100kbps is the number of mismatches or indels per 100kbps when aligning the contigs to the reference genome. A misassembly is an instance where two adjacent stretches of bases in the assembly align either to two very distant or to two highly overlapping stretches of the reference genome. The Quast study defines these metrics in more detail [27].

Assemblies produced from reads corrected with the four programs are very similar according to these measures, with Quake and Lighter yielding the longest contigs and the greatest genome coverage. Surprisingly, the post-correction assemblies have more differences at nucleotide level compared to the pre-correction assemblies, perhaps due to spurious corrections.

#### GAGE human chromosome 14

We also evaluated Lighter’s effect on alignment and assembly using a dataset from the GAGE project [29]. The dataset consists of real 101 × 101 bp paired-end reads covering human chromosome 14 to 35× average coverage (∼36.5M reads). For each error corrector, we tested different *k*-mer sizes (19, 23, 27, 31) and chose the size that yielded the greatest total number of matching aligned nucleotides. For the assembly experiment, we set each error corrector’s *k*-mer size to match that selected in the alignment experiment. Also for each assembly experiment, we tested different *k*-mer sizes for Velvet (47, 53, 57, 63, 67) and chose the one that yielded the greatest NG50.

Error correction’s effect on Bowtie 2 alignment statistics are shown in Table 5. We used Bowtie 2 with default parameters to align the reads to an index of the human chromosome 14 sequence of the hg19 build of the human genome. As before, Lighter yields the greatest improvement in fraction of reads aligned, whereas Quake and BLESS yield the greatest improvement in fraction of aligned bases that match the reference, with Lighter very close behind.

**Table 5.** Alignment statistics for the GAGE chromosome 14 dataset

|  | *k* | Read Level |  | Base Level |  |
| --- | --- | --- | --- | --- | --- |
|  |  | Mapped Reads | Increase (%) | Matches/aligned base (%) | Increase (%) |
| Original | - | 35,993,147 | - | 98.507 | - |
| Quake | 19 | 32,547,091 | -9.57 | 99.845 | 1.36 |
| SOAPec | 19 | 36,116,405 | 0.34 | 98.768 | 0.26 |
| Musket | 19 | 36,316,699 | 0.90 | 99.109 | 0.61 |
| BLESS | 27 | 36,301,816 | 0.86 | 99.411 | 0.92 |
| Lighter | 19 | 36,320,688 | 0.91 | 99.235 | 0.74 |

We repeated this experiment using BWA-MEM as the aligner instead of Bowtie 2 (Supplementary Table 6) and found that the tools performed similarly.

We also tested error correction’s effect on de novo assembly of this dataset using Velvet for assembly and Quast to evaluate the quality of the assembly. For each tool we tested different *k*-mer sizes (19, 23, 27, 31) and chose the one that yielded the greatest NG50. Results are shown in Table 6. Overall, Lighter’s accuracy on real data is comparable to other error correction tools, with Lighter and BLESS achieving the greatest N50, NG50 and coverage.

**Table 6.** De novo assembly statistics for the GAGE chromosome 14 dataset

|  | N50 | NG50 | Edits / 100kbp | Misassemblies | Coverage (%) |
| --- | --- | --- | --- | --- | --- |
| Original | 5290 | 3861 | 139.46 | 1263 | 78.778 |
| Quake | 4829 | 3520 | 141.59 | 1201 | 78.358 |
| SOAPec | 5653 | 4143 | 127.8 | 623 | 79.087 |
| Musket | 5587 | 4105 | 131.17 | 559 | 79.175 |
| BLESS | 5898 | 4345 | 128.4 | 581 | 79.279 |
| Lighter | 5827 | 4280 | 127.69 | 618 | 79.287 |

#### C. elegans

Using the same procedure as in the previous section, we measured the effect of error correction on another large real data using the reads from accession SRR065390. Results are shown in Tables 7 and 8. This run contains real 100 × 100 bp paired-end reads covering the *C. elegans* genome (WBcel235) to 66× average coverage (∼ 67.6M reads). *k*-mer sizes for the error correctors and for Velvet were selected in the same was as for the chromosome 14 experiment. The alignment comparison shows BLESS achieving the greatest increase in fraction of reads aligned, and BLESS and Quake achieving the greatest fraction of aligned bases that match the reference, probably due to their trimming policy. Lighter does the best of the non-trimming tools in the alignment comparison. In the assembly comparison, Lighter and SOAPec achieve the greatest N50, NG50, and coverage.

**Table 7.** Alignment statistics for the *C. elegans* dataset

|  | *k* | Read Level |  | Base Level |  |
| --- | --- | --- | --- | --- | --- |
|  |  | Mapped Reads | Increase (%) | Matches/aligned base (%) | Increase (%) |
| Original | - | 63,017,855 | - | 99.048 | - |
| Quake | 19 | 60,469,150 | -4.04 | 99.834 | 0.79 |
| SOAPec | 19 | 63,032,768 | 0.02 | 99.185 | 0.14 |
| Musket | 23 | 63,060,601 | 0.07 | 99.420 | 0.38 |
| BLESS | 31 | 64,150,807 | 1.80 | 99.744 | 0.70 |
| Lighter | 23 | 63,081,655 | 0.10 | 99.469 | 0.43 |

**Table 8.** De novo assembly statistics for the *C. elegans* dataset

|  | N50 | NG50 | Edits / 100kbp | Misassemblies | Coverage (%) |
| --- | --- | --- | --- | --- | --- |
| Original | 17,330 | 17,317 | 27.66 | 441 | 94.873 |
| Quake | 13,887 | 13,668 | 27.19 | 559 | 94.320 |
| SOAPec | 19,369 | 19,457 | 25.71 | 449 | 95.308 |
| Musket | 18,761 | 18,917 | 28.02 | 438 | 95.288 |
| BLESS | 17,673 | 17,693 | 29.24 | 524 | 94.968 |
| Lighter | 19,222 | 19,333 | 26.9 | 434 | 95.332 |

### Speed, space usage, and scalability

We compared Lighter’s peak memory usage, disk usage, and running time with Quake, Musket and BLESS. These experiments were run on a computer running Red Hat Linux 4.1.2-52 with 48 2.1 GHz AMD Opteron processors and 512G memory. The input datasets are the same simulated *E. coli* datasets with 1% error rate discussed previously, plus the GAGE human chromosome 14 dataset and *C. elegans* dataset.

The measure of space usage is shown in Table 9. BLESS and Lighter achieve constant memory footprint across sequencing depths. While Musket uses less memory than Quake, it uses more than either BLESS or Lighter. BLESS achieves constant memory footprint across sequencing depths, but consumes more disk space for datasets with deeper sequencing. Note that BLESS can be configured to trade off between peak memory footprint and the number of temporary files it creates. Lighter’s algorithm uses no disk space. Lighter’s only sizable data structures are the two Bloom filters, which reside in memory.

**Table 9.**
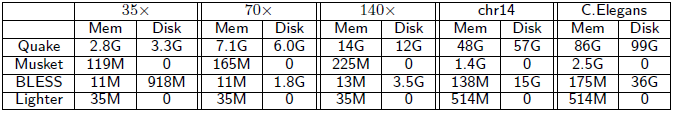
Comparison of four error correction tools based on their memory usage (peak resident memory) and disk usage.

To assess scalability, we also compared running time for Quake, Musket and Lighter using different number of threads. For these experiments we used the simulated *E. coli* dataset with 70× coverage and 1% error. Results are shown in Figure 6. Note that Musket requires at least 2 threads due to its master-slave design. BLESS can only be run with one thread and its running time is 1812s, which is slower than Quake.

**Figure 6.**
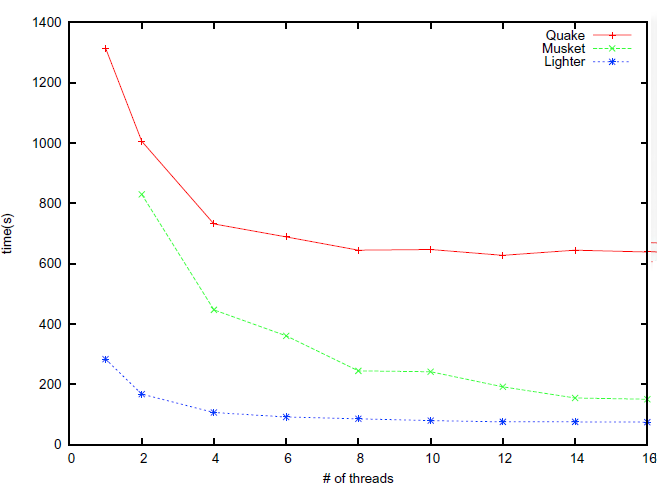
Running times. The running times of Quake, Musket and Lighter on 70× simulated dataset with increasing number of threads

## Discussion

At Lighter’s core is a method for obtaining a set of correct *k*-mers from a large collection of sequencing reads. Unlike previous methods, Lighter does this without counting *k*-mers. By setting its parameters appropriately, its memory usage and accuracy can be held almost constant with respect to depth of sequencing. It is also quite fast and memory-efficient, and requires no temporary disk space.

Though we demonstrate Lighter in the context of sequencing error correction, Lighter’s counting-free approach could be applied in other situations where a collection of solid *k*-mers is desired. For example, one tool for scaling metagenome sequence assembly uses of a Bloom filter populated with solid *k*-mers as a memory-efficient, probabilistic representation of a De Bruijn graph [18]. Other tools use counting Bloom filters [30, 31] or the related CountMin sketch [32] to represent De Bruijn graphs for compression [19] or digital normalization and related tasks [33]. We expect Ideas from Lighter could be useful in reducing the memory footprint of these and other tools.

An important question is how Lighter’s performance can be improved for datasets where coverage is significantly non-uniform, and where solid *k*-mers can therefore have widely varying abundance. In practice, datasets have non-uniform coverage because of ploidy, repeats and sequencing bias. Also, assays such as exome and RNA sequencing intentionally sample non-uniformly from the genome. Even in standard whole-genome DNA sequencing of a diploid individual, *k*-mers overlapping heterozygous variants will be about half as abundant as *k*-mers overlapping only homozygous variants. Lighter’s ability to classify the heterozygous *k*-mers deteriorates as a result, as shown in the section “Effect of ploidy on Bloom Filter B” above. Hammer [11] relaxes the uniformity-of-coverage assumption and favors corrections that increase the multiplicity of a *k*-mer, without using a threshold to separate solid from non-solid *k*-mers. A question for future work is whether something similar can be accomplished in Lighter’s non-counting regime, or whether some counting (e.g. with a counting Bloom filter [30, 31] or CountMin sketch [32]) is necessary.

A related issue is systematically biased sequencing errors, i.e. errors that correlate with the sequence context. One study demonstrates this bias in data from the Illumina GA II sequencer [34]. This bias boosts the multiplicity of some incorrect *k*-mers, causing problems for error correction tools. For Lighter, increased multiplicity of incorrect *k*-mers causes them to appear more often (and spuriously) in Bloom filters A and/or B, ultimately decreasing accuracy. It has also been shown that these errors (a) tend to have low base quality, and (b) tend to occur only on one strand or the other [34]. Lighter’s policy of using a 5th-percentile threshold to classify low-quality positions as untrusted will help in some cases. However, because Lighter canonicalizes *k*-mers (as do many other error correctors), it loses information about whether an error tends to occur on one strand or the other.

Lighter has three parameters the user must specify: the *k*-mer length *k*, the genome length *G*, and the subsampling fraction *α*. While the performance of Lighter is not overly sensitive to these parameters (see Figures 3 and 5), it is not desirable to leave these settings to the user. In the future, we plan to extend Lighter to estimate *G*, along with appropriate values for *k*, and *α*, from the input reads. This could be accomplished with methods proposed in the KmerGenie [35] and KmerStream [21] studies.

Lighter is free open source software released under the GNU GPL license, and has been compiled and tested on Linux, Mac OS X and Windows computers. The software and its source are available from https://github.com/mourisl/Lighter/.

## Author’s contributions

LS and BL designed and analyzed the method. LS implemented the software. LS, LF and BL evaluated the software and wrote the manuscript.

## Acknowledgements

The authors thank Jeff Leek for helpful discussions. *Funding:* National Science Foundation grant ABI-1159078 to LF and a Sloan Research Fellowship to BL.

